# Control of human pancreatic beta cell kinome by GLP-1R biased agonism

**DOI:** 10.1101/2023.02.25.530040

**Authors:** Jiannan Xiao, Liliane El Eid, Teresa Buenaventura, Raphaël Boutry, Amélie Bonnefond, Ben Jones, Guy A Rutter, Philippe Froguel, Alejandra Tomas

## Abstract

**Aim:** To determine the kinase activity profiles of human pancreatic beta cells downstream of GLP-1R balanced *versus* biased agonist stimulations.

**Materials and methods:** This study analysed the kinomic profiles of human EndoC-βh1 cells following vehicle and glucagon-like peptide-1 receptor (GLP-1R) stimulation with the pharmacological agonist exendin-4, as well as exendin-4-based biased derivatives exendin-phe1 and exendin-asp3 for acute (10-minute) *versus* sustained (120-minute) responses, using PamChip® protein tyrosine kinase (PTK) and serine/threonine kinase (STK) assays. The raw data were filtered and normalised using BioNavigator. The kinase analyses were conducted with R, mainly including kinase-substrate mapping and Kyoto Encyclopedia of Genes and Genomes (KEGG) pathway analysis.

**Results:** The present analysis reveals that kinomic responses are distinct for acute *versus* sustained GLP-1R agonist (GLP-1RA) exposure, with individual responses associated with agonists presenting specific bias profiles. According to pathway analysis, several kinases, including JNKs, PKCs, INSR and LKB1, are important GLP-1R signalling mediators, constituting potential targets for further research on biased GLP-1R downstream signalling.

**Conclusion:** Results from this study suggest that differentially biased exendin-phe1 and exendin-asp3 can modulate distinct kinase interaction networks. Further understanding of these mechanisms will have important implications for the selection of appropriate anti-T2D therapies with optimised downstream kinomic profiles.

## Introduction

Type 2 diabetes (T2D) is the commonest form of diabetes mellitus, arising due to defective insulin production and insulin resistance (1). Therefore, increasing insulin secretion from pancreatic beta cells to compensate for insulin resistance and maintain blood glucose levels is an important principle of T2D therapies (2). Insulin secretion is a key physiological process involving the co-operation of multiple signalling pathways (3). In particular, glucagon-like peptide-1 receptor (GLP-1R) targeting to potentiate this process is highly effective for the treatment of T2D, with GLP-1R agonists (GLP-1RAs) such as Liraglutide, Exenatide and Semaglutide already in use successfully in the clinic (4). There are however associated deleterious effects of GLP-1RA therapies linked to gastrointestinal disturbances (5) and, given its global prevalence, there remains a huge unmet need for more efficacious agents for T2D treatment. It is therefore necessary to identify possible directions to improve and prolong the beneficial effects of GLP-1RAs while reducing the negative ones (6).

In the context of deepening understanding of multidimensional networks in G protein-coupled receptor (GPCR) signal transduction, the concept of “biased agonism”, which describes ligand selectivity for specific downstream pathways, has emerged (7). For the GLP-1R, the G_α_s subunit is the main G protein subtype binding to active receptors, triggering adenylate cyclase (AC) activation, and resulting in the production of cyclic adenosine monophosphate (cAMP) (8). Additionally, some studies highlight G protein-independent signalling elicited by the recruitment of β-arrestins to active GLP-1Rs (9). Single amino acid changes in the reference GLP-1RA exendin-4 (Exenatide) were found to result in pronounced GLP-1R biased signalling (10). Specifically, two exendin-4 derivatives, exendin-phe1 and exendin-asp3, were associated with preferential G_α_s or β-arrestin recruitment, respectively. GLP-1R activation with these biased compounds was also linked to changes in receptor trafficking profiles, with exendin-phe1 triggering slow GLP-1R internalisation followed by rapid recycling, while exendin-asp3 is associated with rapid GLP-1R internalisation and preferential sorting of the receptor for lysosomal degradation. As a result, GLP-1R desensitisation was reduced and insulin release prolonged with exendin-phe1 in rat INS-1 832/3 beta cells, and *in vivo* glucoregulation improved in mice (10). GLP-1R biased agonists are therefore potential attractive options to improve existing incretin therapies (11). Indeed, the newly approved incretin mimetic Tirzepatide may depend partly on biased GLP-1R agonism for its high therapeutic efficacy (12). However, the complex molecular mechanisms leading to the divergent responses of biased GLP-1RAs remain unclear.

In this context, phosphorylation-regulated signalling pathways play an essential role in enabling cells to respond quickly and effectively to various cellular signals and stressors (13). Protein kinases are an important group of intracellular enzymes involved in the regulation of cellular functions such as proliferation, apoptosis, and metabolism (14). In eukaryotic cells, serine/threonine (Ser/Thr) and tyrosine (Tyr) are the main amino acid phosphosites (15). Analysis of cellular “kinomic” profiles is a recent field of study whereby global kinase signalling responses are determined. This approach has been chiefly applied into cancer research, resulting in the development of specific kinase inhibitors used successfully in cancer therapy (16). However, despite its potential to increase our understanding of the complex mechanisms linked to GLP-1R signalling, kinase network analysis has not been previously applied to pancreatic beta cells in the context of GLP-1RA stimulation (17), or for the analysis of downstream effects of biased GLP-1R signalling.

In this report, we describe the acute (10-minute) *versus* long term (120-minute) protein tyrosine kinase (PTK) and serine/threonine kinase (STΚ) activity profiles of human EndoC-βH1 pancreatic beta cells under the effects of balanced (exendin-4) *versus* oppositely biased (exendin-phe1, exendin-asp3) GLP-1RAs using PamChip® technology. The major interest of this study is the identification of critical kinases activated by GLP-1R downstream signalling, including the determination of those differentially regulated by biased agonism.

## Materials and Methods

### Peptide agonists

Peptide agonists (exendin-4, exendin-phe1, exendin-asp3) were purchased from by Wuxi Apptec at >90% purity.

### Cell culture and treatments

EndoC-βh1 cells, obtained under licence from Human Cell Design, were cultured in DMEM containing 5.6 mM D-glucose (Sigma-Aldrich), supplemented with 2% bovine serum albumin (BSA) fraction V (Sigma-Aldrich), 50 μM β-mercaptoethanol (Sigma-Aldrich), 10 mM nicotinamide, 5.5 μg/ml transferrin, 6.7 ng/ml sodium selenite and 1% penicillin/streptomycin. The culture plates were coated with DMEM 4500 mg/L glucose (D6546 Sigma), 10% Fetal Bovine Serum (FBS), 1% penicillin/streptavidin, 2 μg/mL fibronectin and 1% extracellular matrix (ECM) (18) prior to cell plating. Cells were starved in 3 mM glucose media overnight prior to pre-incubation for 1 hour in KREBS buffer (140 mM NaCl, 3.6 mM KCl, 1.5 mM CaCl_2_, 0.5 mM MgSO_4_, 0.5 mM NaH_2_PO_4_, 2 mM NaHCO_3_, 10 mM HEPES, 1% BSA, saturated with 95% O_2_ / 5% CO_2_; pH 7.4) containing 0.5 mM glucose before stimulation in 15 mM glucose KREBS buffer with and without addition of 100 nM exendin-4, exendin-phe1 or exendin-asp3 for the indicated times. Immediately following stimulation, cells were lysed with 1% Nonidet-P40 in TNE buffer [20 mM Tris, 150 mM NaCl, 1 mM ethylenediaminetetraacetic acid (EDTA), pH 7.4], supplemented with protease and phosphatase inhibitor cocktails.

### PTK activity profiling

PTK profiles were determined using the PTK PamChip® Array, a flow-through microarray assay which contains 196 unique peptide sequences consisting of 13-15 amino acids with putative endogenous phosphorylation sites that are substrates of Tyr kinases. The principle of the assay is to determine the phosphorylation of the substrate peptide by detection of fluorescence following binding to FITC-conjugated PY20 anti-phosphotyrosine antibody. PTK mixtures were prepared using standard protocols provided by Pamgene. Sample incubation, detection and preliminary analysis were performed in the PamStation 12 machine according to the manufacturer’s instructions. The microarray analysis was run for 94 cycles, while the CCD camera of the PamStation 12 recorded images at kinetic read cycles 32–93 and at end level read cycle at 10, 20, 50, 100, and 200 msec (17). Instrument manipulation as initial sample/array processing and image capture was performed using Evolve (Pamgene) software. In the final step, data was normalized with the BioNavigator analysis software tool (Pamgene) and the mean fluorescence signal intensity changes [log_2_(FC)] of four replicates per experimental group was obtained and normalised to the corresponding vehicle control.

### STK activity profiling

STK profiles were determined using the STK PamChip® Array, a flow-through microarray assay which contains 144 unique peptide sequences which, as for the PTK array, consist of 13-15 amino acids containing putative endogenous phosphorylation sites that are substrates of Ser/Thr kinases. The phosphorylation of these substrate peptides was detected by fluorescence using a two-step assay (17) according to standard protocols provided by Pamgene. The STK mixtures contain both primary antibodies and a FITC-labelled secondary antibody. Sample incubation, detection and preliminary analysis were performed in the PamStation 12 according to the manufacturer’s instructions. The microarray analysis was run for 124 cycles, while the CCD camera of the PamStation 12 recorded images at kinetic read cycles 92-132 and read the final cycle at 10, 20, 50, 100, and 200 msec (17). Instrument manipulation as initial sample/array processing and image capture was performed using Evolve (Pamgene) software. In the final step, data were normalized with the BioNavigator analysis software tool (Pamgene) and as for the PTK analysis, the mean fluorescence signal intensity changes [log_2_(FC)] of four replicates per experimental group were obtained and normalised to the corresponding vehicle control.

### Matching substrates to upstream kinases

Kinase-substrate information was obtained from PhosphoNET, Uniprot and PhosphoSite databases. The top 50 Ser/Thr kinases and 25 Tyr kinases with prediction V2 scores higher than 300 were selected for each phosphosite from PhosphoNET (Supplementary Files 1 and 2). Kinase-substrate mapping was performed in R.

#### Upstream kinase analysis

We modified the kinase-substrate enrichment analysis (KSEA) formula (19) to use the average phosphosite log_2_(FC) weighted by the prediction V2 scores for each phosphosite of a given kinase. The modified z kinase score formula is as follows:

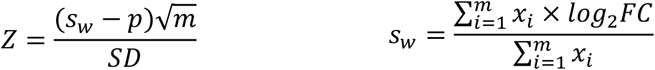

Where s_w_ represents the weighted mean log_2_(FC) of the substrate subset for a given kinase, p denotes the mean log_2_(FC) of all phosphosites in the raw dataset, m is the number of phosphosite substrates identified for each kinase, SD represents the standard deviation of the log_2_(FC) across all phosphosites and x represents the PhosphoNET predictor V2 score. The scores were assumed to be normally distributed, therefore the p-values were determined by one-tail test to estimate the probability that the real scores were at least as extreme as the measured scores. Data sorting, analysis and calculation were performed in Excel and R. Kinase scores were visualized in R mainly using the “pheatmap” package (v1.0.12, https://github.com/raivokolde/pheatmap).

### Pathway enrichment analysis

To conduct a representative analysis of kinase lists, the “enrichKEGG” function in R package ClusterProfiler v4.0 was applied to determine whether the kinases identified are enriched within specific pathways (20). Statistical thresholds of p-value < 0.05 and q-value < 0.05 were chosen to select significantly enriched KEGG terms and pathways. A dot plot of the top 10 KEGG terms and pathways, ranked by kinase ratio and coloured by adjusted p-value, was depicted. A Venn diagram was generated to identify the kinases shared within these pathways using Venn webtools (https://bioinformatics.psb.ugent.be/webtools/Venn/), with a list of kinases illustrated.

### RNA-seq database analysis

High-throughput sequencing data were retrieved from Gene Expression Omnibus (GEO) databases to assess the level of expression of specific kinase genes. Gene expression data from EndoC-βH1 cell line was obtained using GEO accession GSM3333914. Additionally, raw read count values of 18 non-diabetic and 39 T2D islet samples from GSE164416 series were analysed using DESeq2 (21) in order to compare differentially expressed kinase genes, followed by integration with our kinomic analysis results.

### Generation of kinase tree maps and kinase networks

An annotated phylogenetic kinase tree was generated using the KinMap online tool (Beta version, http://kinhub.org/kinmap/) (22). Data source was selected as “Disease associated CTTV”, and specific Ser/Thr and Tyr kinases highlighted in the kinase tree. Kinases associated with “diabetes mellitus” and “metabolic disease” were visualised for comparison and reference. The interaction network of our candidates was visualised with String version 11.0 (https://string-db.org). Additionally, KEGG and reactome pathway analyses were presented as STRING networks.

### Other statistical analyses

Other statistical analyses were performed in Excel, R (version 4.1.3), and Prism 8 (GraphPad Software, San Diego USA). Data visualisation was performed in R, Prism 8 and BioRender (https://biorender.com/). Unless otherwise stated, p-values < 0.05 were considered statistically significant.

### Supporting information

All the source code from the present study can be found at github: Kinomic-assay.

## Results

### Kinomic effects of acute versus sustained GLP-1RA stimulation in human EndoC-βh1 pancreatic beta cells

The kinomic assays were performed following the workflow summarized in Figure 1 to analyse active kinases for two treatment duration groups: a) 10-minute (acute), and b) 120-minute (sustained), in EndoC-βh1 cells, a cell line that we have previously shown capable of displaying both cAMP and insulin secretion responses to GLP-1R stimulation (23). We chose to assess acute *versus* sustained effects as differences in glucoregulatory responses between biased GLP-1RAs are exacerbated over time (10). For each group, four treatment conditions were established: vehicle, exendin-4, exendin-phe1 and exendin-asp3 (at 100nM each). Figure 2A presents the peptide sequences for each of the three GLP-1RAs tested in the study. EndoC-βh1 cell lysates were collected immediately for each condition and kept at -80°C until further analysis. Equal amounts of lysates were applied for PTK and STK assays on peptide microarray chips (PamChip®). PamGene read the signal intensity of peptide phosphorylation from raw data images and completed primary quality control steps. The list of substrates included in the assay is shown in Supplementary File 3. Temporally regulated, agonist-specific kinase activity profiles triggered by GLP-1R activation were detected in EndoC-βh1 cells, based on the differential signal intensities of phosphorylated peptides measured for each experimental condition (Supplementary Figure 1). Although substrate phosphorylation indicated the existence of differences between the conditions, it was not clear which was the pattern of agonist-induced kinase activity for each GLP-1RA tested without further analysis. To assess this, we modified the KSEA formula (24) to consider the affinity of individual kinases towards binding to specific substrates. In this way, kinase scores were calculated to determine the degree of specific downstream kinase activity changes following GLP-1R stimulation. Here, we present a list of kinases that showed significant activity changes compared to vehicle exposure under at least one of the conditions tested (Figure 2B, C). All the identified kinases were confirmed to be expressed in EndoC-βh1 cells at the mRNA level [GSE118588, (25)]. Consistent with our substrate phosphorylation results, more pronounced changes were detected in Tyr kinase activity after sustained *versus* acute GLP-1RA exposure (Figure 2B). For STKs, responses were more variable and presented marked changes over vehicle conditions in both acute and sustained experimental groups (Figure 2C). Importantly, the close kinase scores obtained downstream of acute exendin-4 and exendin-asp3 stimulation indicated that the two agonists lead to similar STK signalling profiles in EndoC-βh1 cells at an early time point (Figure 2C). Additionally, differences detected in acute *versus* sustained kinase action for these two agonists compared to exendin-phe1 suggest a different mode of action for the latter, and may provide clues to identify underlying pathways leading to the previously described differences in acute *versus* sustained glucoregulatory effects of this biased GLP-1RA (10) (26).

**Figure 1.**
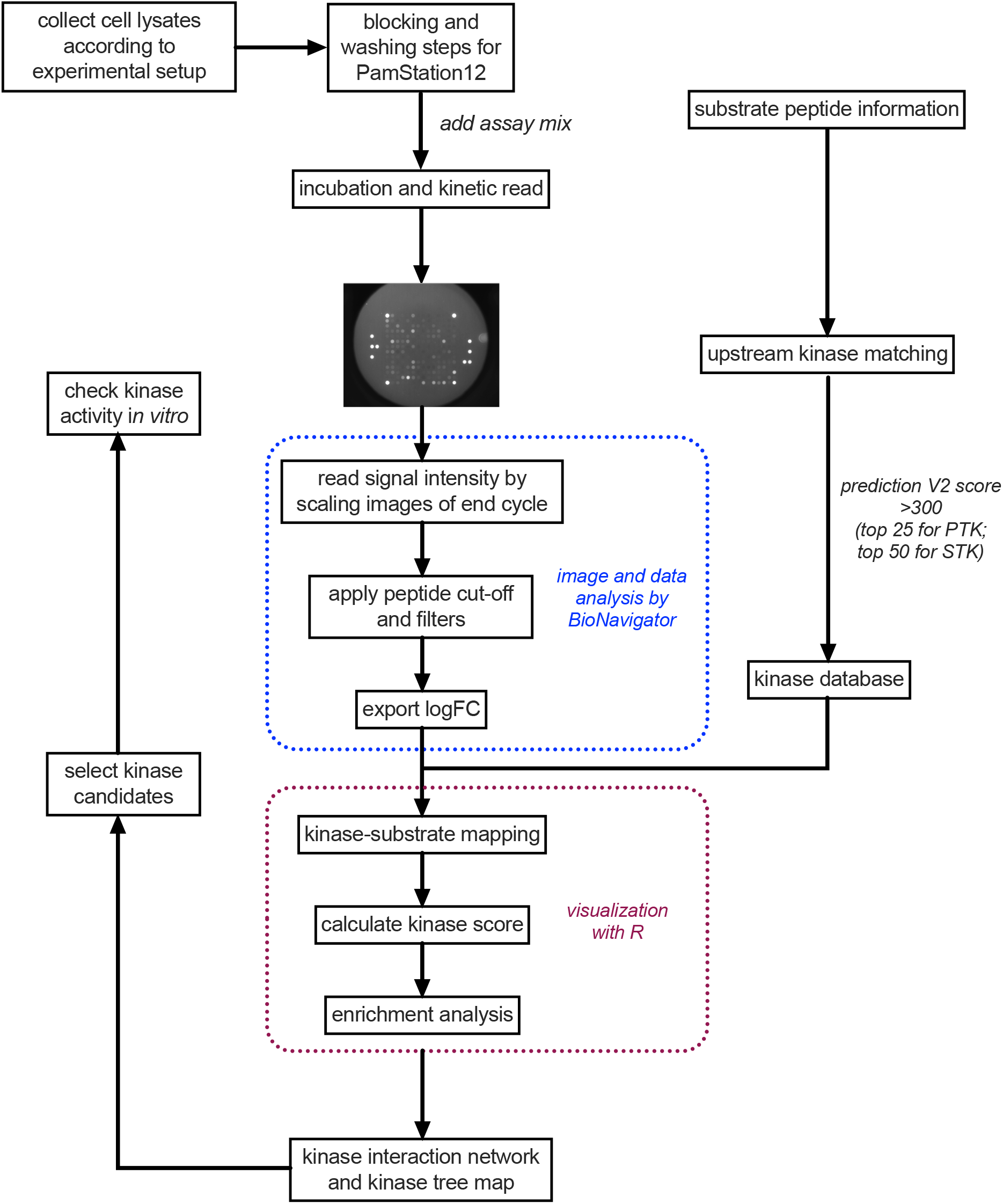
Schematic of kinomic analysis workflow.

**Figure 2.**
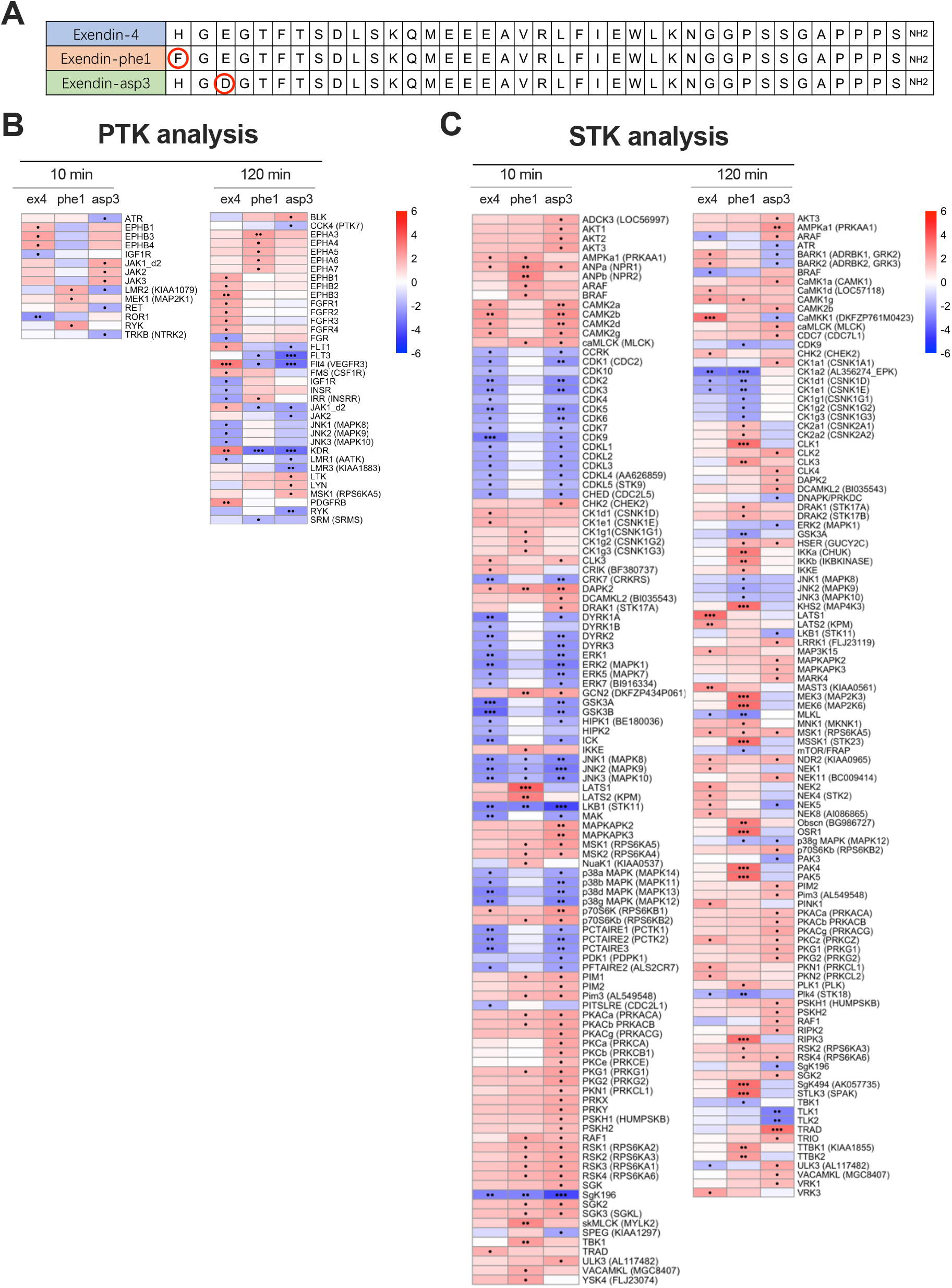
Kinase activity profiles of EndoC-βh1 cells following acute and sustained GLP-1R stimulations with exendin-4, -phe1 and -asp3. (**A**) Peptide sequences of exendin-4 (ex4), exendin-phe1 (phe1), and exendin-asp3 (asp3). (**B**) Heatmap visualization of PTK kinase z scores, predicting Tyr kinase activity downstream of the indicated treatment. (**C**) As for (B) for STK scores. Positive z scores (shown in red) indicate increased kinase activity, while negative z scores (shown in blue) indicate decreased kinase activity compared to vehicle conditions; •p<0.05, ••p<0.01, •••p<0.001; p-values determined by one-tail t-tests.

### Pathway analysis reveals key kinases involved in GLP-1R downstream signalling

To further understand the impact of changes in individual kinase activity on global beta cell kinase networks, we performed pathway enrichment analysis for those kinases showing significant activity changes downstream of GLP-1RA stimulations. The top 10 relevant KEGG pathways were sorted by kinase ratio, as shown in Figure 3A, B; we focused on the first three pathways from each kinomic dataset. The MAPK signalling pathway was highlighted as the most enriched pathway for both PTK and STK analyses, in agreement with previous reports linking MAPK family activities to pancreatic beta cell function (27). For instance, PKA-activated MAPKs play a crucial role in accelerating the cell cycle, which is associated with maintenance of beta cell mass (28). According to the Venn diagram analysis of the kinases implicated (Figure 3C, D), there are 13 Tyr kinases and 15 Ser/Thr kinases common to all our top selected KEGG pathways. For the Ser/Thr kinases, those identified belong mainly to the Rapidly Accelerated Fibrosarcoma (RAF) and the MAPK/ERK families, both previously shown to play pivotal roles downstream of GLP-1R activation (29).

**Figure 3.**
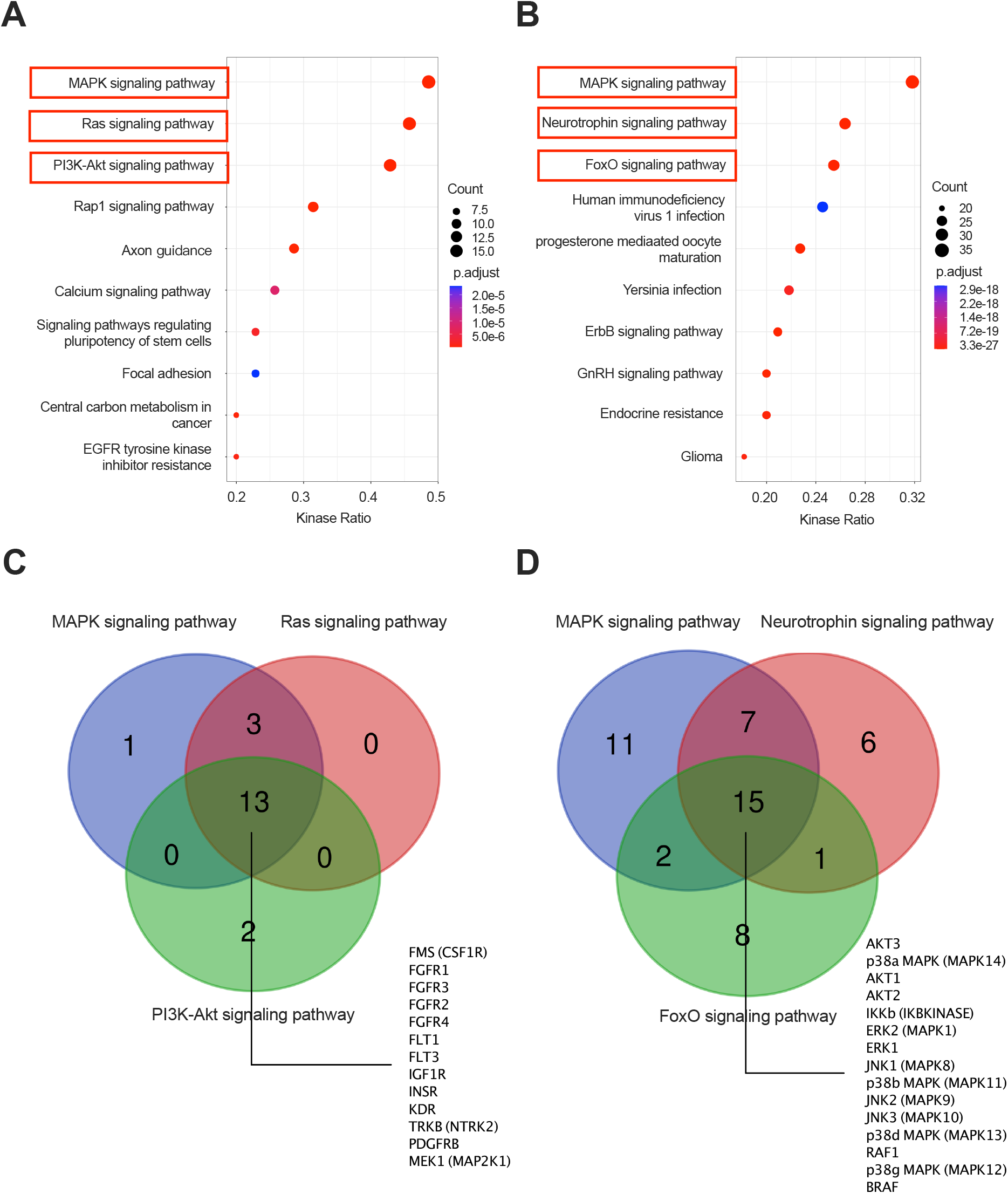
Pathway analysis for predicted kinases. (**A**) Top 10 KEGG pathways for identified Tyr kinases ranked by kinase ratio. (**B**) As for (A) for Ser/Thr kinases. Dot sizes reflect kinase counts and dot colours adjusted p-values; Red boxes represent selected pathways. (**C**) Venn diagram of selected KEGG pathways for Tyr kinases. (**D**) As for (C) for Ser/Thr kinases. Numbers shown in the overlapping areas indicate intersecting kinases involved in the represented pathways.

### Biased agonist-mediated changes in downstream kinase signatures

Besides comparing kinase activity changes caused by acute and sustained GLP-1R stimulations, we also focused here on the identification of specific kinase responses downstream of GLP-1R biased agonist signalling. To this end, we assessed kinase activities downstream of differentially biased agonists exendin-phe1 and exendin-asp3 by recalculating fold changes in substrate phosphorylation levels for the biased agonists *versus* exendin-4. In this way, we obtained new kinase scores that more directly represent the differences in kinase activity triggered by exendin-phe1 and exendin-asp3 relative to the parental compound, and we show here a list of kinases with predicted activity differences under at least one experimental condition (Figure 4A, B). Interestingly, our results suggest that there are some kinases being differentially regulated by exendin-phe1 compared to exendin-asp3 stimulation. For example, for PTKs, acute stimulation with exendin-phe1 leads to increased activity of Lemur tyrosine kinase 2 (LMR2), while exendin-asp3 stimulation for the same time causes a reduction in LMR2 activity compared to exendin-4 (Figure 4A). Moreover, although the change trends in Tyr kinase activity were similar for both biased agonists following sustained exposure, the effect of exendin-phe1 was more pronounced than that of exendin-asp3 at this time point for several kinases, including cJun N-terminal kinases (JNKs) and insulin receptor (INSR) (Figure 4A). Results for STKs showed more dramatic differences between both biased agonists, especially after sustained treatment, with prolonged exendin-phe1 responses again leading to significantly increased effects compared to exendin-asp3 (Figure 4B). Examples of differentially activated kinases include cyclin-dependent kinases (CDKs) and dual-specificity tyrosine-regulated kinases (DYRKs), which showed increased activity following acute exendin-phe1 stimulation, while PKCs showed higher acute activation with exendin-asp3 (Figure 4B). Additionally, MAPK kinases (MAP2K/MEKs: MEK3, MEK6) and p21-activated kinases (PAKs: PAK4, PAK5) showed greater discrepancies between exendin-phe1 and exendin-asp3 at the sustained time-point. According to a further KEGG analysis, the Ras, MAPK and PI3K-Akt signalling pathways were again the top 3 enriched pathways related to Tyr kinases (Supplementary Figure 2A). However, only the MAPK signalling pathway remained as one of the three most enriched for Ser/Thr kinases, this being the only one from the top three clearly related to pancreatic beta cell function (Supplementary Figure 2B). To provide more detailed evidence for the regulation of the pathways involved, a systems biology approach was conducted and depicted using STRING v11.0 (https://string-db.org) to analyse interactions amongst the three top-related kinases pathways for Tyr kinases (Figure 4C) and within the MAPK signalling pathway for Ser/Thr kinases (Figure 4D). These analyses showed that the selected pathways were highly overlapped and correlated, although some kinases may act independently from others. Specifically, most PTKs can be linked to JAK2, FGFR2 and JAK3 (Figure 4C), while BRAF and PRKCB were the most centred amongst the STKs (Figure 4D), suggesting that they may play important roles in regulating beta cell biased GLP-1RA responses.

**Figure 4.**
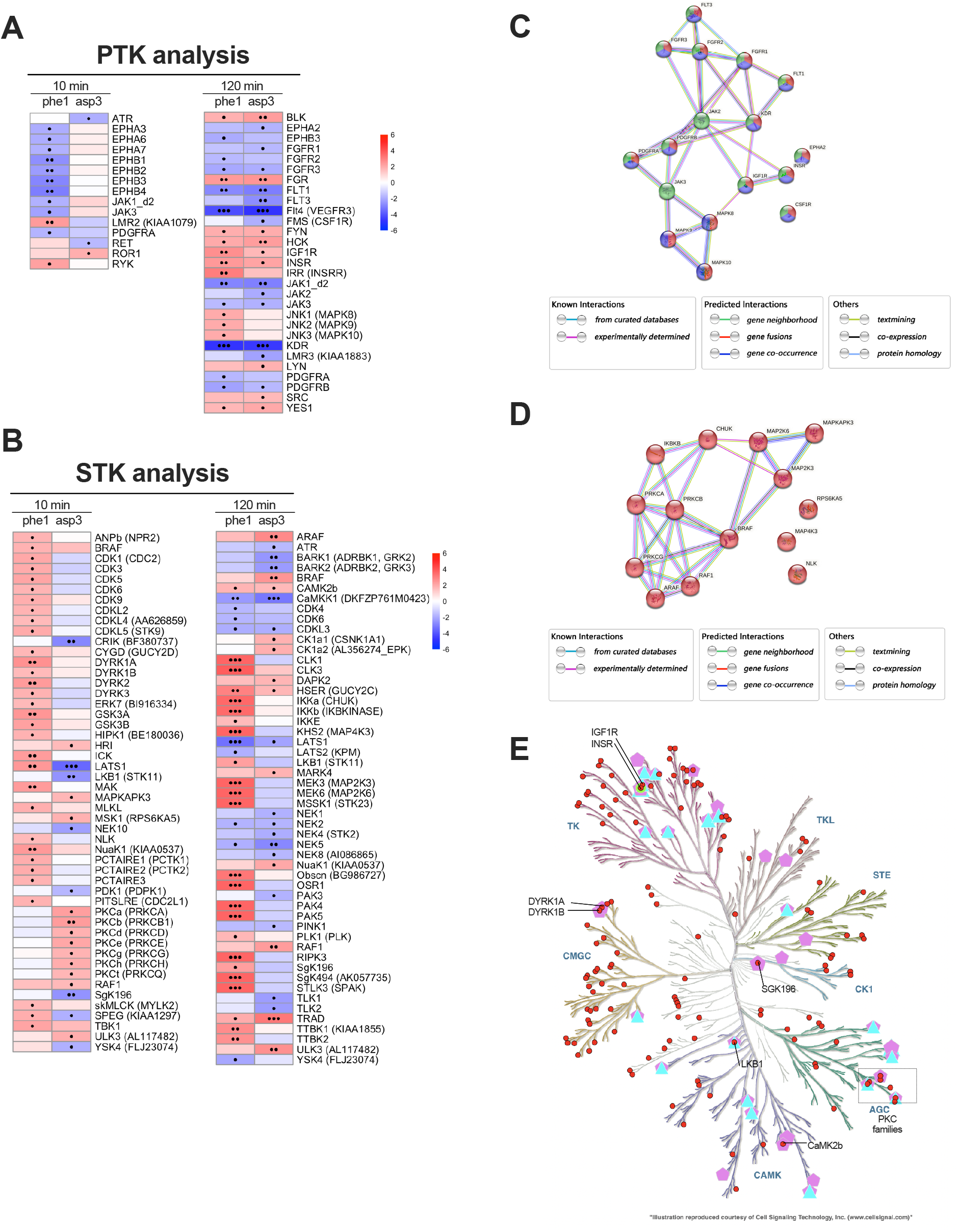
Biased agonist *versus* exendin-4 kinase activity profiles. (**A**) Heatmap of recalculated PTK z scores for exendin-phe1 (phe1) and exendin-asp3 (asp3), indicating Tyr kinase activity changes *versus* exendin-4. (**B**) As for (A) for Ser/Thr kinase activity changes compared to exendin-4. Positive z scores (shown in red) indicate increased, while negative z scores (shown in blue) indicate decreased kinase activity; •p<0.05, ••p<0.01, •••p<0.001; p-values were determined by one-tail t-test. (**C**) Kinase interaction network for Tyr kinases involved in the pathways selected; red: Ras pathway; blue: MAPK pathway; green: PI3K-Akt pathway; (**D**) Kinase interaction network for Ser/Thr kinases involved in the MAPK signalling pathway. A medium confidence (minimum required interaction score of 0.004) was selected for depicted interactions; filled nodes represent proteins with known or predicted 3D structures, while empty nodes represent proteins of unknown 3D structure; node colours represent distinct pathways. (**E**) Kinase tree map: red circles correspond to kinases shown in Figure 4A, B; triangles are kinases related to diabetes mellitus; green circles are kinases related to abnormal glucose homeostasis; pentagons are kinases related to metabolic disease.

As pancreatic beta cells constitute more than 50% of the mass of human islets (30), we additionally confirmed expression of the kinases assessed in this study in a human islet RNA-seq database that contained 18 non-diabetic and 39 T2D islets [GSE164416, (31)] (Supplementary Figure 3). Interestingly, nearly all the kinases showing significant changes in mRNA expression between healthy and T2D islets were also included in the list of differentially regulated kinases by GLP-1R biased signalling (Figure 4A, B). Finally, we classified all human kinases into 8 typical (AGC, CAMK, CK1, CMGC, STE, TK, TKL, Other) and 13 atypical families based on their underlying sequences (22). A kinase map tree was then generated for and annotated with those kinases identified in our study implicated in diabetes mellitus or metabolic disease (Figure 4E). PKCs, INSR, and liver kinase B1 (LKB1), known to be important targets in the development of diabetes, were all suggested to have differential activities under the effects of the biased GLP-1RAs tested. Specifically, PKCs were more active for acute stimulation with exendin-asp3, while LKB1 activity was increased during prolonged treatment with exendin-phe1 (Figure 4B).

## Discussion

The present study has lain a foundation for understanding the effect of GLP-1RAs and biased GLP-1R signalling on the physiology of pancreatic beta cells at the kinomic level. Substrate polypeptides were observed to exhibit differential phosphorylation after acute *versus* sustained treatments with the different GLP-1RAs, indicating that GLP-1R downstream kinase responses are temporally regulated. Corresponding to the substrate results, different kinomic profiles were identified following treatment with GLP-1RAs with opposed bias characteristics, suggesting that careful assessment of these profiles might be a new way to determine the specific effects of signal bias for the GLP-1R. Based on our kinase-substrate mapping analysis, we have unveiled here activity changes linked to multiple Tyr and Ser/Thr kinases. Elucidation of complex pathways requires identification of key active enzymes within these pathways and determination of their roles in the overall regulation of signalling. Notably, our pathway enrichment analysis highlighted the critical role of prominent kinases such as the MAPKs and PKCs in GLP-1R downstream signalling in human beta cells.

Importantly, this is the first study analyzing kinomic profiles of pancreatic beta cells downstream of GLP-1R stimulation. Kinases are activated by phosphorylation, which subsequently activates a cascade of events resulting in the phosphorylation of specific amino acids within substrates (32). Different databases have been designed to recognise kinase-specific phosphosites to facilitate the elucidation of phospho-signalling networks. For example, PhosphoSite database and NetworKIN predictions are recommended for KSEA (19). Here, we used the PhosphoNET database, which predicted more than 300 kinases for our substrates. By weighting the prediction V2 score, we were able to assess the influence of the different kinases in the degree of phosphorylation of each substrate. Although our substrate peptides are relatively well conserved, public databases are mainly generated based for human kinases. Therefore, at this moment this method of analysis may only be suitable for human cell lines such as EndoC-βh1.

Our study particularly aimed to determine signalling differences elicited by biased GLP-1RAs. The KEGG pathway analysis allowed the identification of specific kinase families involved in this process, including: *<u>JNK families</u>*, implicated in the regulation of most cellular processes as one of the main MAPK subfamilies (27). The main functional role of activated JNK is to phosphorylate c-Jun, with Ser63 and Ser73 phosphorylations required for c-Jun transcriptional activation (33). All the JNKs (JNK1,2 and 3) were identified in our kinase analysis as displaying increased activity following exendin-phe1 stimulation over a sustained period. However, JNK inhibition has been suggested to prevent beta cell dysfunction and apoptosis in both human islets and beta cell lines (27). JNK1/2 can mediate interleukin-1β (IL-1β) induced ER Ca^2+^ release, leading to mitochondrial dysfunction in mouse and primary human beta cells (34). Moreover, oxidative stress in pancreatic beta cells under chronic exposure to high glucose levels induce JNK activation, impairing the insulin signalling cascade (35). Conversely, in rat insulin-secreting cells, JNK3 silencing dramatically decreased expression of insulin receptor substrate 2 (IRS2), which regulates beta cell function and proliferation via PI3K-Akt signalling. Importantly, exendin-4 was found to increase JNK3 expression in a previous study on isolated human islets and INS-1E cells (36). It is therefore possible that activation of JNK kinases downstream of GLP-1R action might lead to differential signalling effects than those triggered by chronic hyperglycaemia. Further experimental validation, focusing on phosphorylation of different phosphosites by candidate kinases may be required for in-depth characterization of JNK kinases and their downstream signalling pathways.

Besides JNK, *<u>PKC families</u>* have also been repeatedly identified as a hub in the present kinase network analysis of GLP-1R signalling. More specifically, PKCs have been identified as showing increased activity following acute exendin-asp3 *versus* exendin-phe1 exposure. PKA-independent actions of GLP-1 have previously been shown to depend on PKC, leading to activation of phospholipase C (PLC), elevation of diacylglycerol (DAG), and increased L-type Ca^2+^ and TRPM4/TRPM5 channel activities (37). On the other hand, high glucose induced PKCβ and PKCδ overexpression has been related to GLP-1 resistance (38), with PKCβ inhibition suggested as a potential therapy to ameliorate pancreatic beta cell dysfunction via promoting the degradation of Thioredoxin (TXN)-interacting protein (TXNIP) (39). Moreover, PKCβ is significantly upregulated in islets of T2D patients according to our RNA-seq analysis. It remains to be determined if the favourable profile of exendin-phe1 can be linked to a more nuanced activation of PKC kinases that avoids some of the deleterious effects associated with increased activity of PKCβ and PKCδ.

Additionally, sustained exendin-phe1 exposure also caused higher *<u>INSR</u>* activity. INSR expression is usually high in pancreatic beta cells, but its function in these cells is not yet clear. Downregulation of INSR in INS-1 cells leads to reduced glucose-stimulated insulin secretion (40), potentially linking the effect of exendin-phe1 on INSR with the increased sustained insulin secretion triggered by this compound. However, a recent study suggested that INSR knockout in beta cells results in enhanced insulin secretion (41). Therefore, more evidence is needed to elucidate if INSR signalling downstream of GLP-1R activation is beneficial or detrimental for beta cell function.

Besides the pathways identified in our KEGG analysis, *<u>AMPK signalling</u>* also caught our attention as our analysis points to both LKB1 and CaMKK1 being differentially regulated by acute *versus* prolonged exposure to GLP-1RAs. The LKB1-AMPK pathway regulates beta cell function by controlling glucose coupling by directing glucose-derived carbons into the tricarboxylic acid cycle (TCA) cycle, an important process for the maintenance of mitochondrial structure and function (42). Studying the effect of GLP-1R agonism in the control of this pathway will therefore be particularly important to understand the effects of incretin action on the regulation of beta cell function.

Our analysis of kinomic profiles provides evidence for the importance of factors related to the growth, proliferation, and function of pancreatic beta cells to explain the effects of GLP-1R action. Besides some well-known kinase families, we have identified some kinases whose precise roles in beta cells are still unknown. For example, the DYRK family, which has previously been hinted at being closely related to the control of beta cell function (43), has been shown here to be differentially modulated by biased GLP-1R agonism, with exendin-phe1 being more effective than exendin-asp3 at acute DYRK activation.

So far, we have only performed a computational prediction of kinase activity profiles downstream of GLP-1R stimulation in beta cells, and our results have not yet been individually validated. Future verification of key kinase activity and kinetic profiles by Western blot and FRET-based assays followed by corresponding mechanistic investigations of the functional consequences of modulating specific kinase activities using RNAi or chemical inhibition approaches will be necessary to fully elucidate the kinase signatures of GLP-1R biased *versus* balanced compounds. Furthermore, LC-MS/MS analysis following co-immunoprecipitation, or proximity ligation assays could assist in the identification of specific substrates downstream of the identified kinases. It is worth mentioning that our kinase database would be suitable for the analysis of kinomic profiles from any human cell line using the same substrates. However, as our kinase profiling is based on predicted peptide phosphorylation levels using literature databases, an obvious limitation of the present study is that kinase signalling may be different *in vivo*, as well as in primary islets. In the future, it will be important to find ways to translate our findings on kinase function to *in vivo* and *ex vivo* models of GLP-1R action.

## Conclusion

As the first study to compare kinase profiles of pancreatic beta cells downstream of balanced *versus* biased GLP-1RA stimulations, our analysis has shed new light on our understanding of GLP-1R signalling and biased agonism. We have linked kinase activity profiles to GLP-1R “signal bias” in beta cells, hoping to extract the most central changes from this complex pathway. In particular, we have found that phosphorylation of different amino acid residues within the same kinase may affect kinase function, which could partly explain the dual/controversial role of certain kinases in the control of beta cell function.

## Acknowledgements

This work was supported by MRC grant number MR/R010676/1 to A.T., B.J., G.A.R and P.F., as well as by UKRI COVID-19 Grant Extension Allocation (coA) to A.T. A.T. also acknowledges further support from the EFSD, Diabetes UK, Eli Lilly, the Commonwealth, and the Integrated Biological Imaging Network (IBIN). The authors thank Prof. Amar Abderrahmani for assistance during raw data acquisition.

## References

1. Nauck MA, Wefers J, Meier JJ. Treatment of type 2 diabetes: challenges, hopes, and anticipated successes. Lancet Diabetes Endocrinol. 2021 Aug;9(8):525–44.

2. Taylor SI, Yazdi ZS, Beitelshees AL. Pharmacological treatment of hyperglycemia in type 2 diabetes. J Clin Invest. 2021 Jan 19;131(2):e142243.

3. Campbell JE, Newgard CB. Mechanisms controlling pancreatic islet cell function in insulin secretion. Nat Rev Mol Cell Biol. 2021 Feb;22(2):142–58.

4. Knudsen LB, Lau J. The Discovery and Development of Liraglutide and Semaglutide. Front Endocrinol. 2019 Apr 12;10:155.

5. Lingvay I, Hansen T, Macura S, Marre M, Nauck MA, de la Rosa R, et al. Superior weight loss with once-weekly semaglutide versus other glucagon-like peptide-1 receptor agonists is independent of gastrointestinal adverse events. BMJ Open Diabetes Res Care. 2020 Oct;8(2):e001706.

6. Jones B, McGlone ER, Fang Z, Pickford P, Corrêa IR, Oishi A, et al. Genetic and biased agonist-mediated reductions in β-arrestin recruitment prolong cAMP signaling at glucagon family receptors. J Biol Chem. 2021 Jan;296:100133.

7. Lei S, Clydesdale L, Dai A, Cai X, Feng Y, Yang D, et al. Two distinct domains of the glucagon-like peptide-1 receptor control peptide-mediated biased agonism. J Biol Chem. 2018 Jun;293(24):9370–87.

8. Marzook A, Tomas A, Jones B. The Interplay of Glucagon-Like Peptide-1 Receptor Trafficking and Signalling in Pancreatic Beta Cells. Front Endocrinol. 2021 May 10;12:678055.

9. Diz-Chaves Y, Herrera-Pérez S, González-Matías LC, Lamas JA, Mallo F. Glucagon-Like Peptide-1 (GLP-1) in the Integration of Neural and Endocrine Responses to Stress. Nutrients. 2020 Oct 28;12(11):3304.

10. Jones B, Buenaventura T, Kanda N, Chabosseau P, Owen BM, Scott R, et al. Targeting GLP-1 receptor trafficking to improve agonist efficacy. Nat Commun. 2018 Dec;9(1):1602.

11. Trujillo JM, Nuffer W, Smith BA. GLP-1 receptor agonists: an updated review of head-to-head clinical studies. Ther Adv Endocrinol Metab. 2021 Jan;12:204201882199732.

12. Willard FS, Douros JD, Gabe MBN, Showalter AD, Wainscott DB, Suter TM, et al. Tirzepatide is an imbalanced and biased dual GIP and GLP-1 receptor agonist. JCI Insight. 2020 Sep 3;5(17):e140532.

13. Ardito F, Giuliani M, Perrone D, Troiano G, Muzio LL. The crucial role of protein phosphorylation in cell signaling and its use as targeted therapy (Review). Int J Mol Med. 2017 Aug;40(2):271–80.

14. Attwood MM, Fabbro D, Sokolov AV, Knapp S, Schiöth HB. Trends in kinase drug discovery: targets, indications and inhibitor design. Nat Rev Drug Discov. 2021 Nov;20(11):839–61.

15. Gnad F, Forner F, Zielinska DF, Birney E, Gunawardena J, Mann M. Evolutionary Constraints of Phosphorylation in Eukaryotes, Prokaryotes, and Mitochondria. Mol Cell Proteomics. 2010 Dec;9(12):2642–53.

16. Bhullar KS, Lagarón NO, McGowan EM, Parmar I, Jha A, Hubbard BP, et al. Kinase-targeted cancer therapies: progress, challenges and future directions. Mol Cancer. 2018 Dec;17(1):48.

17. Alack K, Weiss A, Krüger K, Höret M, Schermuly R, Frech T, et al. Profiling of human lymphocytes reveals a specific network of protein kinases modulated by endurance training status. Sci Rep. 2020 Dec;10(1):888.

18. Ravassard P, Hazhouz Y, Pechberty S, Bricout-Neveu E, Armanet M, Czernichow P, et al. A genetically engineered human pancreatic β cell line exhibiting glucose-inducible insulin secretion. J Clin Invest. 2011 Sep 1;121(9):3589–97.

19. Wiredja DD, Koyutürk M, Chance MR. The KSEA App: a web-based tool for kinase activity inference from quantitative phosphoproteomics. Valencia A, editor. Bioinformatics. 2017 Nov 1;33(21):3489–91.

20. Yu G, Wang LG, Han Y, He QY. clusterProfiler: an R Package for Comparing Biological Themes Among Gene Clusters. OMICS J Integr Biol. 2012 May;16(5):284–7.

21. Love MI, Huber W, Anders S. Moderated estimation of fold change and dispersion for RNA-seq data with DESeq2. Genome Biol. 2014 Dec;15(12):550.

22. Eid S, Turk S, Volkamer A, Rippmann F, Fulle S. KinMap: a web-based tool for interactive navigation through human kinome data. BMC Bioinformatics. 2017 Dec;18(1):16.

23. Buenaventura T, Kanda N, Douzenis PC, Jones B, Bloom SR, Chabosseau P, et al. A Targeted RNAi Screen Identifies Endocytic Trafficking Factors That Control GLP-1 Receptor Signaling in Pancreatic β-Cells. Diabetes. 2018 Mar 1;67(3):385–99.

24. Casado P, Rodriguez-Prados JC, Cosulich SC, Guichard S, Vanhaesebroeck B, Joel S, et al. Kinase-Substrate Enrichment Analysis Provides Insights into the Heterogeneity of Signaling Pathway Activation in Leukemia Cells. Sci Signal [Internet]. 2013 Mar 26 [cited 2022 May 15];6(268). Available from: https://www.science.org/doi/10.1126/scisignal.2003573

25. Lawlor N, Márquez EJ, Orchard P, Narisu N, Shamim MS, Thibodeau A, et al. Multiomic Profiling Identifies cis-Regulatory Networks Underlying Human Pancreatic β Cell Identity and Function. Cell Rep. 2019 Jan;26(3):788-801.e6.

26. Lucey M, Pickford P, Bitsi S, Minnion J, Ungewiss J, Schoeneberg K, et al. Disconnect between signalling potency and in vivo efficacy of pharmacokinetically optimised biased glucagon-like peptide-1 receptor agonists. Mol Metab. 2020 Jul;37:100991.

27. Kassouf T, Sumara G. Impact of Conventional and Atypical MAPKs on the Development of Metabolic Diseases. Biomolecules. 2020 Aug 29;10(9):1256.

28. Jiang Y, Zhu L, Wu D, Ni Y, Huang C, Ye H, et al. Type IIB PKA is highly expressed in β cells and controls cell proliferation via regulating Cyclin D1 expression. FEBS J. 2022;289(10):2865–76.

29. Sidarala V, Kowluru A. The Regulatory Roles of Mitogen-Activated Protein Kinase (MAPK) Pathways in Health and Diabetes: Lessons Learned from the Pancreatic β-Cell. Recent Pat Endocr Metab Immune Drug Discov. 2017;10(2):76–84.

30. Campbell JE, Newgard CB. Mechanisms controlling pancreatic islet cell function in insulin secretion. Nat Rev Mol Cell Biol. 2021 Feb;22(2):142–58.

31. Wigger L, Barovic M, Brunner AD, Marzetta F, Schöniger E, Mehl F, et al. Multi-omics profiling of living human pancreatic islet donors reveals heterogeneous beta cell trajectories towards type 2 diabetes. Nat Metab. 2021 Jul;3(7):1017–31.

32. Bilbrough T, Piemontese E, Seitz O. Dissecting the role of protein phosphorylation: a chemical biology toolbox. Chem Soc Rev. 2022;51(13):5691–730.

33. Li L, Feng Z, Porter AG. JNK-dependent Phosphorylation of c-Jun on Serine 63 Mediates Nitric Oxide-induced Apoptosis of Neuroblastoma Cells. J Biol Chem. 2004 Feb;279(6):4058–65.

34. Verma G, Bhatia H, Datta M. JNK1/2 regulates ER–mitochondrial Ca 2+ cross-talk during IL-1β–mediated cell death in RINm5F and human primary β-cells. Newmeyer DD, editor. Mol Biol Cell. 2013 Jun 15;24(12):2058–71.

35. Kawamori D, Kaneto H, Nakatani Y, Matsuoka T aki, Matsuhisa M, Hori M, et al. The Forkhead Transcription Factor Foxo1 Bridges the JNK Pathway and the Transcription Factor PDX-1 through Its Intracellular Translocation. J Biol Chem. 2006 Jan;281(2):1091–8.

36. Nakano R, Nakayama T, Sugiya H. Biological Properties of JNK3 and Its Function in Neurons, Astrocytes, Pancreatic β-Cells and Cardiovascular Cells. Cells. 2020 Jul 29;9(8):1802.

37. Shigeto M, Cha CY, Rorsman P, Kaku K. A role of PLC/PKC-dependent pathway in GLP-1-stimulated insulin secretion. J Mol Med. 2017 Apr;95(4):361–8.

38. Pan X, Chen J, Wang T, Zhang M, Wang H, Gao H. Essential Role Of High Glucose-Induced Overexpression Of PKCβ And PKCδ In GLP-1 Resistance In Rodent Cardiomyocytes. Diabetes Metab Syndr Obes Targets Ther. 2019 Nov;Volume 12:2289–302.

39. He S, Wan Y, Li L, Tang X, Wu W, Liu S, et al. PKCβ Inhibition Promotes TXNIP Degradation to Ameliorate Pancreatic β-Cell Dysfunction. Pharmacology. 2022 Jul 21;1–17.

40. Wang J, Gu W, Chen C. Knocking down Insulin Receptor in Pancreatic Beta Cell lines with Lentiviral-Small Hairpin RNA Reduces Glucose-Stimulated Insulin Secretion via Decreasing the Gene Expression of Insulin, GLUT2 and Pdx1. Int J Mol Sci. 2018 Mar 26;19(4):985.

41. Skovsø S, Panzhinskiy E, Kolic J, Cen HH, Dionne DA, Dai XQ, et al. Beta-cell specific Insr deletion promotes insulin hypersecretion and improves glucose tolerance prior to global insulin resistance. Nat Commun. 2022 Feb;13(1):735.

42. Rourke JL, Hu Q, Screaton RA. AMPK and Friends: Central Regulators of β Cell Biology. Trends Endocrinol Metab. 2018 Feb;29(2):111–22.

43. Rachdi L, Kariyawasam D, Aïello V, Herault Y, Janel N, Delabar JM, et al. Dyrk1A induces pancreatic β cell mass expansion and improves glucose tolerance. Cell Cycle. 2014 Jul 15;13(14):2221–9.

