## Supplementary Figures for "Control of human pancreatic beta cell kinome by GLP-1R biased agonism"

**Supplemental Files**

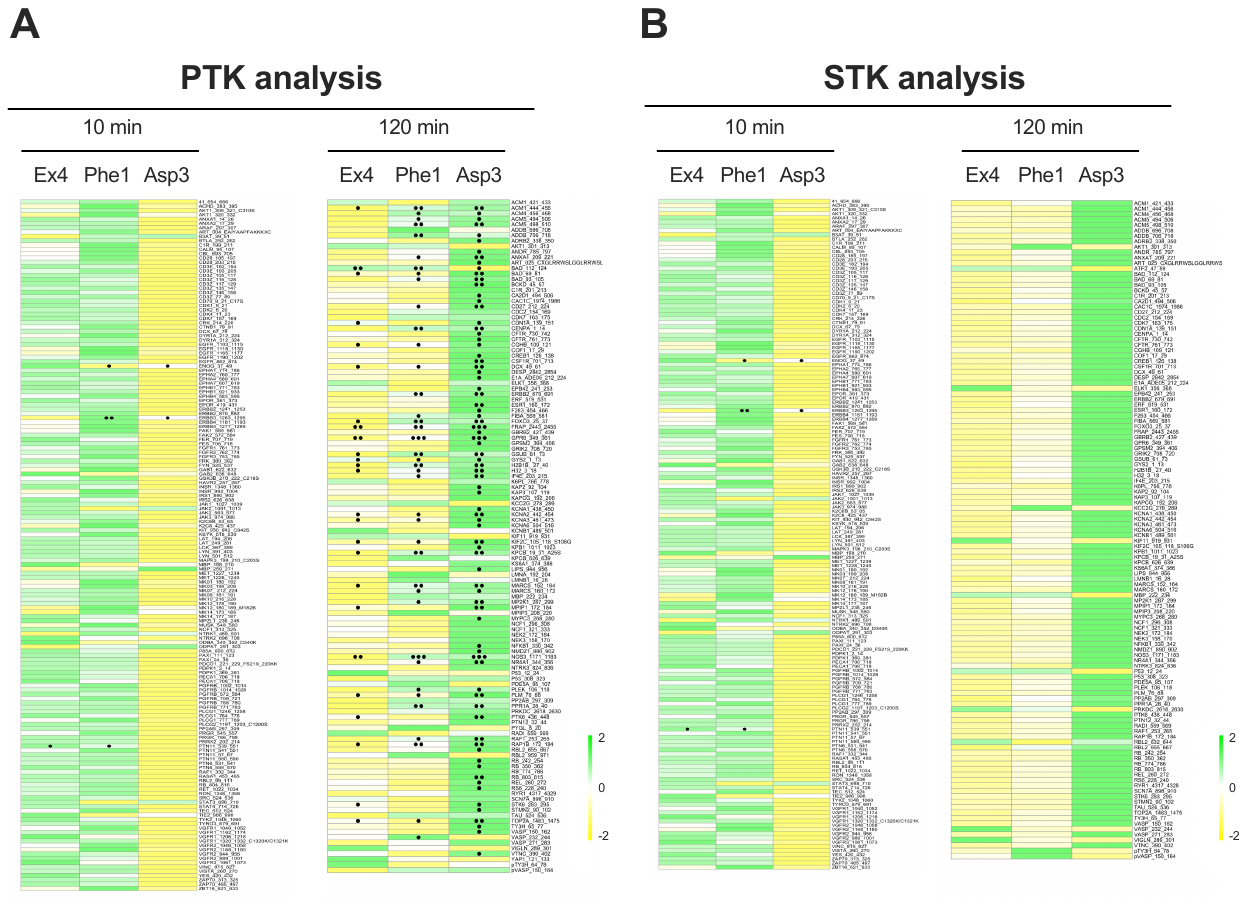

**Supplementary Figure 1.** **PTK and STK substrate phosphorylation levels.** (**A**) Heatmap of PTK substrate phosphorylation levels, displaying the log-transformed signal intensity fold changes in Tyr substrate phosphorylation over vehicle conditions for all quality-controlled peptides included in the assay. (**B**) As for (A) for STK substrate phosphorylation levels; •p<0.05, ••p<0.01, •••p<0.001.

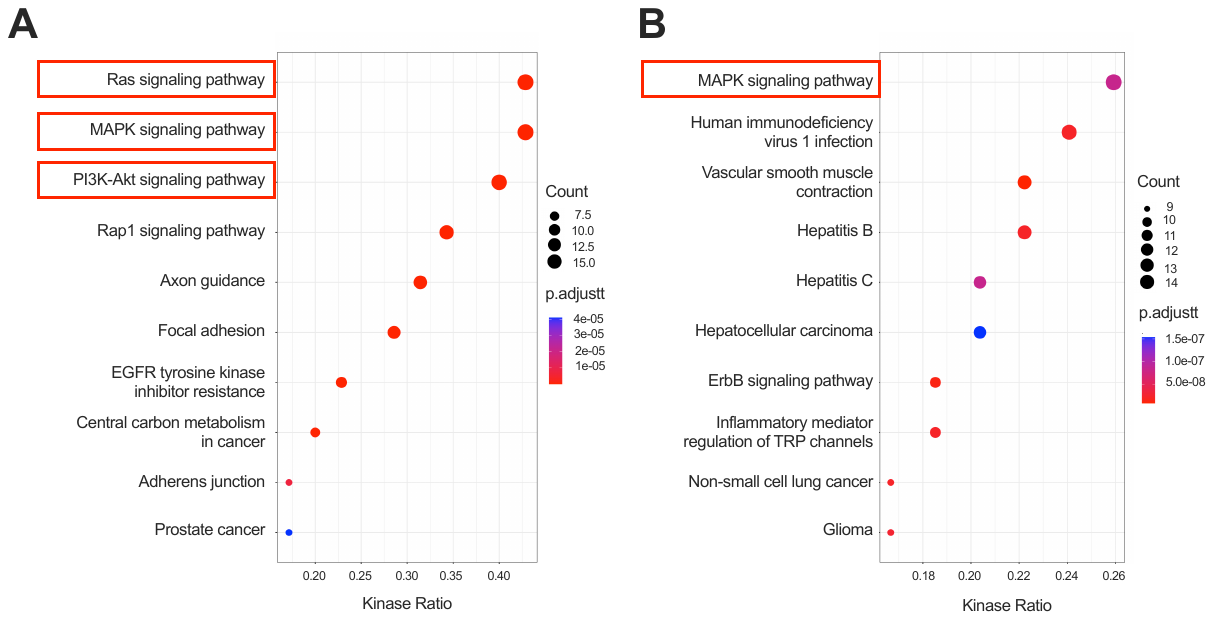

**Supplementary Figure 2. Pathway analysis for candidate kinases for differential biased agonist effects.** (**A**) Top 10 KEGG pathways of enriched Tyr kinases ranked by kinase ratio. (**B**) As for (A) for Ser/Thr kinases; dot sizes reflect kinase counts and dot colours reflect adjusted p-values; red boxes represent selected pathways.

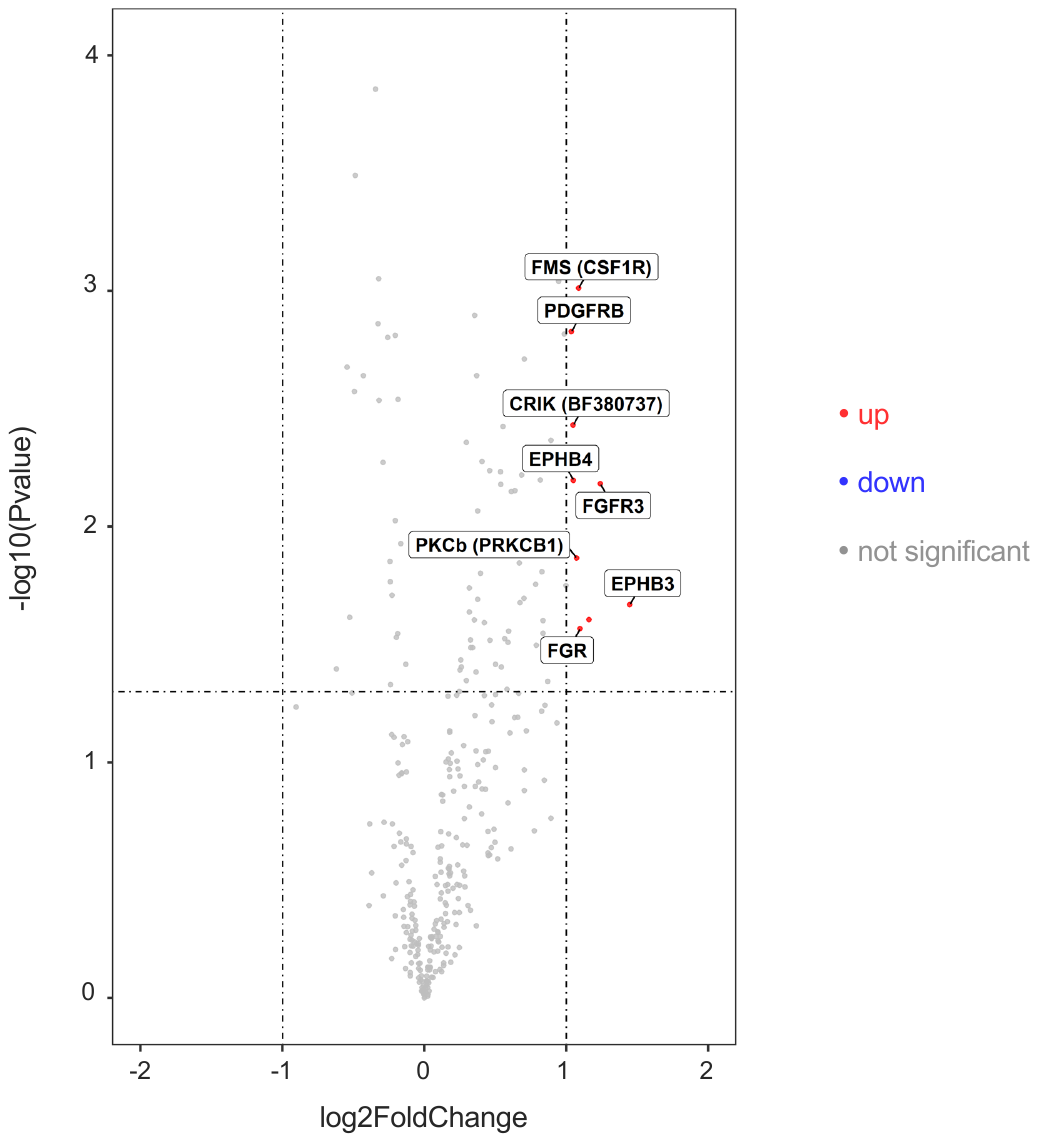

**Supplementary Figure 3. Volcano plots depicting kinase gene expression levels from RNA sequencing datasets of T2D *versus* non-diabetic islets.** Results were considered statistically significant with p<0.05, and log_2_(FC) > 1; red dot indicate upregulated mRNA levels in T2D islets; blue dots indicate downregulated mRNA levels in T2D islets; grey dots indicate non-significant changes in kinase gene expression.

**Supplementary File 1.** ptkdataset1, containing top 25 PTKs.

**Supplementary File 2.** stkdataset1, containing top 50 STKs.

**Supplementary File 3.** List of substrates.
